# CD8+ T-cell-mediated immunoediting influences genomic evolution and immune evasion in murine gliomas

**DOI:** 10.1101/705152

**Authors:** J. Robert Kane, Junfei Zhao, Takashi Tsujiuchi, Brice Laffleur, Aayushi Mahajan, Ganesh Rao, Angeliki Mela, Crismita Dmello, Víctor A. Arrieta, Li Chen, Daniel Y. Zhang, Edgar Gonzalez-Buendia, Catalina Lee-Chang, Ting Xiao, Gerson Rothschild, Uttiya Basu, Craig Horbinski, Maciej S. Lesniak, Amy B. Heimberger, Raul Rabadan, Peter D. Canoll, Adam M. Sonabend

## Abstract

Cancer immunoediting shapes tumor progression by the selection of tumor cell variants that can evade immune recognition. Given the immune evasion and intra-tumor heterogeneity intrinsic to gliomas, we hypothesized that CD8+ T-cells mediate immunoediting in these tumors. We evaluated glioma progression in the absence of CD8+ T-cells by depleting this immune cell population in transgenic murine gliomas. Upon transplantation, gliomas that developed in the absence of CD8+ T-cells engrafted poorly in recipients with intact immunity but engrafted well in those with CD8+ T-cell depletion. Gliomas developed in absence of CD8+ T-cells exhibited increased chromosomal instability, MAPK signaling, gene fusions, and macrophage/microglial infiltration. MAPK activation correlated with macrophage/microglial recruitment in this model and in the human disease. Our results indicate that CD8+ T-cells mediate immunoediting during gliomagenesis, influencing the genomic stability of glioma and its microenvironment, leading to immune evasion.

**Significance:** Immune evasion renders cancer resistant to anti-tumoral immunity. Therapeutic intervention often fails for gliomas because of the plasticity of tumor cell variants that resist immune surveillance. Our results demonstrate a mechanism of immune evasion in gliomas that derives from CD8+ T-cells during the development and progression of this disease.

## Introduction

Glioblastoma (GBM) is the most common and malignant brain cancer in adults with a median overall survival of approximately 21 months [1]. This tumor develops multiple mechanisms of immune suppression to include T-cell anergy, exhaustion, negative regulation, apoptosis, and sequestration in the bone marrow [2, 3]. As a consequence, the efficacy of immunotherapy is hindered.

Cancer immunoediting has been proposed as a mechanism where the adaptive and innate immunity sculpts the immunogenicity of developing tumors, selecting for tumor cell variants that can evade this immune surveillance [4, 5]. This understudied mechanism provides partial explanation as to why many tumors do not respond to various immunotherapies. If anti-tumoral immunity is capable of recognizing some tumor cell variants and not others, immunoediting likely influences the evolutionary path of the cancer. Immunoediting relies on tumor cell heterogeneity— a hallmark of GBM [6]. We recently described the disappearance of immunogenic glioma cell variants following PD-1 blockade in recurrent GBM patients who responded to treatment [7]. This suggests that immunoediting may occur not only during formation of these gliomas, but also in the setting of immunotherapy. In spite of these and other clinical observations that suggest immunoediting in gliomas, this phenomenon has not been experimentally investigated in this disease.

We hypothesized that CD8+ T-cells subject gliomas to immunoediting. By depleting this immune cell population during development of transgenic murine gliomas, we investigated how these immune effectors influence tumor immunogenicity and progression. Upon transplantation, gliomas developed in the absence of CD8+ T-cells engrafted poorly in recipients with intact immunity, but readily grafted in hosts with CD8+ T-cell depletion, implying a more immunogenic profile. Moreover, these gliomas exhibited increased activation of MAPK signaling, mitoses, aneuploidy, gene fusions, and macrophage/microglial infiltration. Our results reveal that CD8+ T-cells elicit a selection pressure that drives glioma progression towards an immune evasive phenotype, while simultaneously protecting genomic integrity during the evolution of cancer.

## Results

### Gliomas that develop in the absence of CD8+ T-cells are more immunogenic yet exhibit distinct oncogenic features

We investigated the effects of CD8+ T-cells on glioma development using a transgenic PDGF^+^*Pten*^-/-^ murine glioma model. In this model, gliomas are induced *de novo* by retroviral vector-based expression of Cre, which leads to loss of the *Pten* tumor suppressor gene (in Pten^lox/lox^ mice) and over-expression of PDGF. Resultant gliomas accumulate genetic alterations that resemble those of the human disease [8]. CD8+ T-cell-mediated immunoediting would require glioma infiltration by these lymphocytes. In order to investigate this hypothesis, we evaluated the presence of CD4+ and CD8+ T-cells at different stages of glioma progression. At 21 days post-virus injection (d.p.i) for glioma induction, which is an early phase of glioma development that precedes the accumulation of the genetic alterations and histological features of high-grade gliomas [8], there was both CD4+ and CD8+ T-cell infiltration in the glioma (**Supplementary Fig. S1A**). We then evaluated T-cell infiltration at 35 d.p.i., a period in which this model exhibits the genetic alterations and histology of high-grade gliomas [8]. Given that these gliomas exhibit considerable growth within weeks, we normalized T-cell infiltration by the average glioma size (estimated by bioluminescence) for each time point (**Supplementary Fig. S1B/C**). This analysis revealed a decline in the fraction of CD4+ as well as CD8+ T-cells in the glioma at 35 d.p.i. compared to that exhibited at 21 d.p.i. (p=0.0189 [CD4+], p=0.0026 [CD8+]; **Supplementary Fig. S1D**).

To investigate the effects of CD8+ T-cells on glioma development, we depleted this immune cell population starting one week prior to retroviral injection, and throughout glioma development. To investigate this effect on glioma immunogenicity, we generated gliomas in hosts with CD8+ T-cell depletion and subsequently transplanted these glioma cells into recipients with or without CD8+ T-cell depletion (**Fig. 1A**). Depletion was achieved by administration of an anti-CD8 antibody as demonstrated by flow cytometry of splenocytes (**Fig. 1B**).The absence of CD8+ T-cells during glioma development had no effect on glioma growth, as mice treated with the anti-CD8 antibody had comparable survival to mice treated with the isotype control antibody (Log-rank p=0.7563; **Fig. 1C**). This is consistent with similar results in a transgenic PDGFB^+^RCAS^-^Stat3^-/-^ glioma model, in which CD8+ T-cells had no effect on overall survival in tumor-bearing mice [9]. Implantation of glioma cells that originated in mice with no CD8+ T-cells into recipients that also had depletion of CD8+ T-cells led to development of gliomas in all cases, with a median survival of 24 days post-glioma induction due to glioma burden (**Fig. 1D)**. In contrast, implantation of the same glioma cells into recipients with intact immunity (treated with the isotype control antibody) led to engraftment in only 1 out of 4 mice, with the remaining animals surviving past 220 days, without any evidence of glioma or neurological symptoms (Log-rank p=0.0180). Known chemokines and cytokines released during the elimination phase of cancer immunoediting [4] were increased in the gliomas that developed in the presence of CD8+ T-cells relative to the CD8+ T-cell depleted background (ANOVA p=0.0106, **Fig. 1E**).

**Figure 1:**
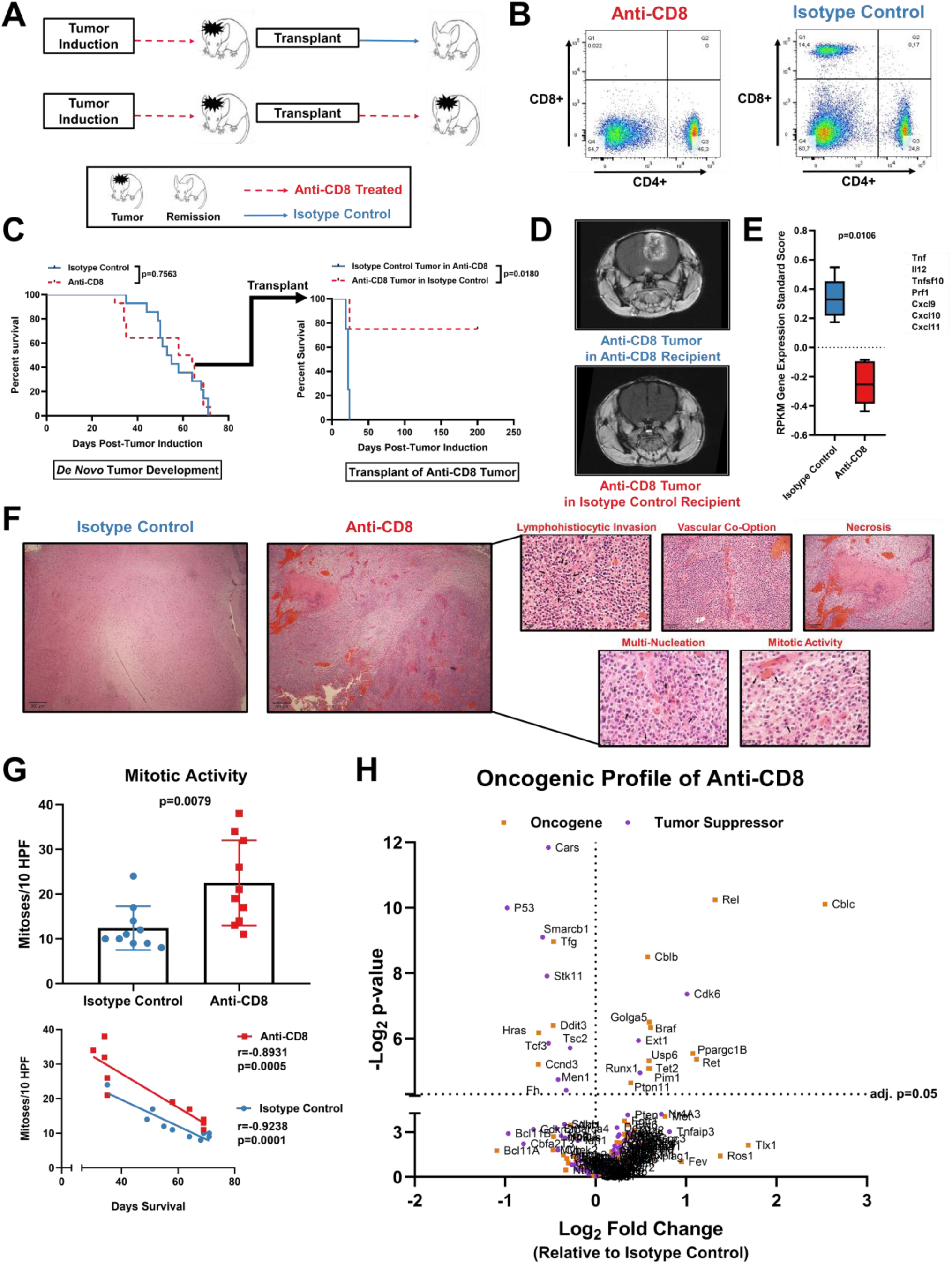
Gliomas that develop in the absence of CD8+ T-cells are more immunogenic yet display increased oncogenic features. (A) Scheme of experimental design to investigate glioma immunoediting. (B) Representative flow cytometry plots demonstrate CD8+ T-cell depletion by administration of Anti-CD8 antibody, but not by isotype control antibody. (C) Survival analysis of animals treated with anti-CD8 and isotype control antibodies prior to and during tumor formation on (n=28). Survival analysis of animals that received the transplant of gliomas treated with anti-CD8 and isotype control antibodies (n=8). (D) Representative MRI of mice implanted with gliomas generated in the absence of CD8+ T-cells. (E) Known signals of the cancer immunoediting elimination are shown by RPKM gene expression standardized score between animals treated with anti-CD8 and isotype control antibodies (ANOVA p=0.0106). Histopathological evaluation by H&E staining in tumors from mice treated with isotype control and anti-CD8 antibodies (F). Tumors from mice treated with the anti-CD8 antibody possessed increased histopathological features of malignancy as shown by representative micrographs. (G) Mitotic figures identified per 10 high-powered fields (HPFs) compared between tumors treated with the anti-CD8 and isotype control antibody (n=20). (H) Relative expression (RNA-seq) of oncogenes and tumor suppressors. Volcano plots show all known oncogenes and tumor suppressors as plotted by their RPKM log_2_ fold change versus adjusted p-value by differential expression analysis in each group.

Gliomas that developed in the absence of CD8+ T-cells exhibited a distinct histology relative to those that developed in the presence of CD8+ T-cells (**Fig. 1F**). In the former group, H&E staining revealed increased lymphohistiocytic infiltrates in addition to a higher extent of necrosis. Vasculature in these gliomas was co-opted by neoplastic cells that were dysmorphic and multi-nucleated. We observed more mitotic activity in gliomas that developed in the absence of CD8+ T-cells (mean mitoses per high power field (HPF): 22.5) as compared to gliomas that developed in the presence of CD8+ T-cells (mean mitoses per HPF: 12.4; p=0.0079; **Fig. 1G**). Mitotic activity showed an inverse correlation with survival (r=-0.8931, p=0.0005 [anti-CD8]; r=-0.9238, p=0.0001 [isotype control]).

Known oncogenes and glioma suppressor genes were examined by their relative expression between gliomas that developed in the presence versus absence of CD8+ T-cells. We found that the gliomas that developed in the absence of CD8+ T-cells exhibited differential expression of both oncogenes and tumor suppressors relative to gliomas that developed in the setting of intact immunity (**Fig. 1H**).

### Gliomas that develop in the absence of CD8+ T-cells exhibit an increased frequency of chromosomal deletions and gene fusions

We investigated whether the absence of CD8+ T-cells during glioma development has an effect on the tumor genome. We performed paired somatic and germline exome sequencing as well as paired-end RNA-seq of gliomas that developed in the presence or absence of CD8+ T-cells. Overall, there were only a few non-synonymous mutations detected, mostly in TP53, consistent with our previous characterization of this glioma model [8]. There was no change in the number of point mutations between these gliomas (p=0.5922; **Fig. 2A**). Yet gene fusions occurred more frequently in the gliomas that developed in the absence of CD8+ T-cells. Specifically, the mean fusion count of gliomas that developed in the absence of CD8+ T-cells was 2.5 versus 0.5 in the presence of CD8+ T-cells (p=0.0309; **Fig. 2B**). In the group of gliomas that developed in the absence of CD8+ T-cells, one glioma showed two neoantigens derived from two separate gene fusion events with predicted high binding affinity to MHC-1 (H2-K^b^ and H2-D^b^).

**Figure 2:**
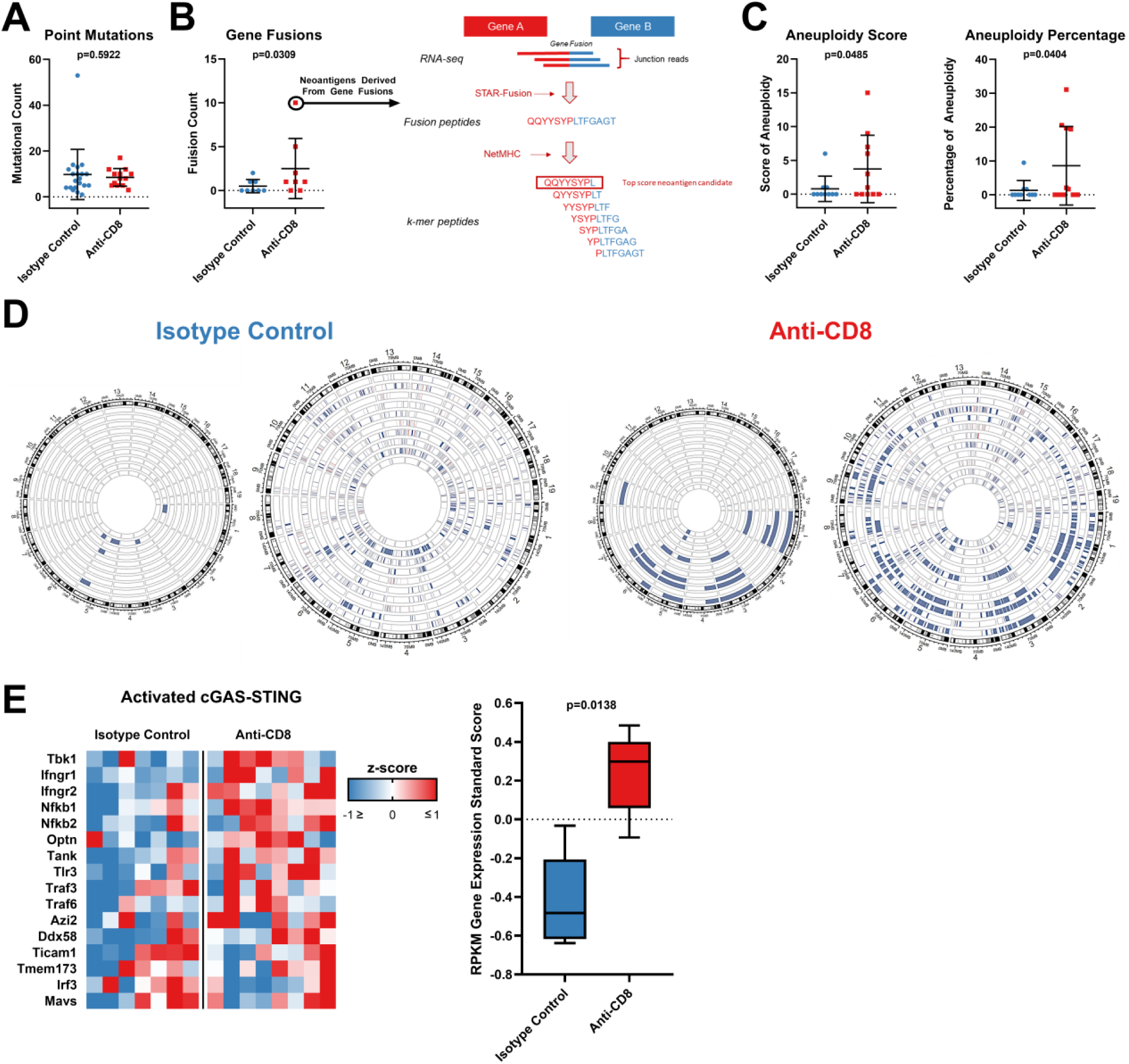
Loss of CD8+ T-cells increases genomic instability in gliomas. (A) Number of point mutations (n=33) and (B) gene fusions (n=16) compared between tumors from animals treated with anti-CD8 and isotype control antibodies. Predicted neoantigens derived from gene fusions from H2-K^b^ and H2-D^b^ are shown via schematic representation of the neoantigen prediction pipeline. (C) Aneuploidy score (n=21) and percentage of aneuploidy across the genome (n=21) compared between tumors from animals treated with anti-CD8 and isotype control antibodies. (D) Chromosomal loss compared between tumors from animals treated with anti-CD8 and isotype control antibodies (n=21; p=0.034). Each wheel represents a track of a tumor for the copy number variation analysis. Blue represents deletion while red represents amplification. The left schematic of each panel depicts only deep copy number variations while the right also depicts shallow variations. (E) The RPKM z-score of genes involved in cGAS-STING signaling is compared between tumors developed in the absence of CD8+ T-cells and the isotype control (n=15; 0.5176 greater mean standard score in tumors developed in the absence of CD8+ T-cells; ANOVA p=0.0138).

Gliomas that developed in the absence of CD8+ T-cells also exhibited increased aneuploidy relative to those that developed in the presence of CD8+ T-cells. Aneuploidy was defined by somatic copy number alterations [10]. The mean aneuploidy score of gliomas that developed in the absence of CD8+ T-cells was 3.7 while it was 0.8 in the presence of CD8+ T-cells (p=0.0485; **Fig. 2C**). We next used an aneuploidy score calculation to infer copy number alterations evenly across the genome as previously described [11]. This analysis revealed that gliomas that developed in the absence of CD8+ T-cells had aneuploidy in 8.587% of the genome while that of the isotype control was only in 1.281% (p=0.0404).

Copy number variation events were mapped in the genome. The increase in aneuploidy exhibited by gliomas that developed in the absence of CD8+ T-cells primarily consisted of chromosomal losses (p=0.034; **Fig. 2D**). As chromosomal instability has been reported to drive innate immune activation via cGAS-STING signaling [12], we evaluated the gene expression associated with cGAS-STING (*Tbk1*) activation by RNA-seq [13]. This analysis revealed increased expression of *Tbk1* and evidence-based associated genes in gliomas that developed in the absence of CD8+ T-cells (ANOVA p=0.0138; **Fig. 2E**). Our results indicate that CD8+ T-cells protect against aneuploidy and gene fusions in these gliomas, whereas these genomic alterations are associated with the activation of the STING pathway.

### ERK and p38 signaling is activated in gliomas that develop in the absence of CD8+ T-cells

Serine/threonine kinase *Braf* as well as phosphatase *Ptpn11* are upstream activators of the MAPK signaling pathway that have been described to have oncogenic properties [14]. Both genes are involved in the activation of the ERK cascade of the MAPK signaling pathway. We recently described the over-representation of activating mutations in these genes in recurrent GBM patients who responded to PD-1 blockade [7]. Given that these mutations were associated with susceptibility to immunotherapy, we hypothesized that murine gliomas that developed in the absence of CD8+ T-cells may exhibit similar alterations. Murine gliomas did not have mutations in these genes, yet RNA-seq analysis revealed that both *Braf* (p=0.0232) and *Ptpn11* (p=0.0256) were over-expressed in gliomas that developed in the absence of CD8+ T-cells (**Fig. 3A**).

**Figure 3:**
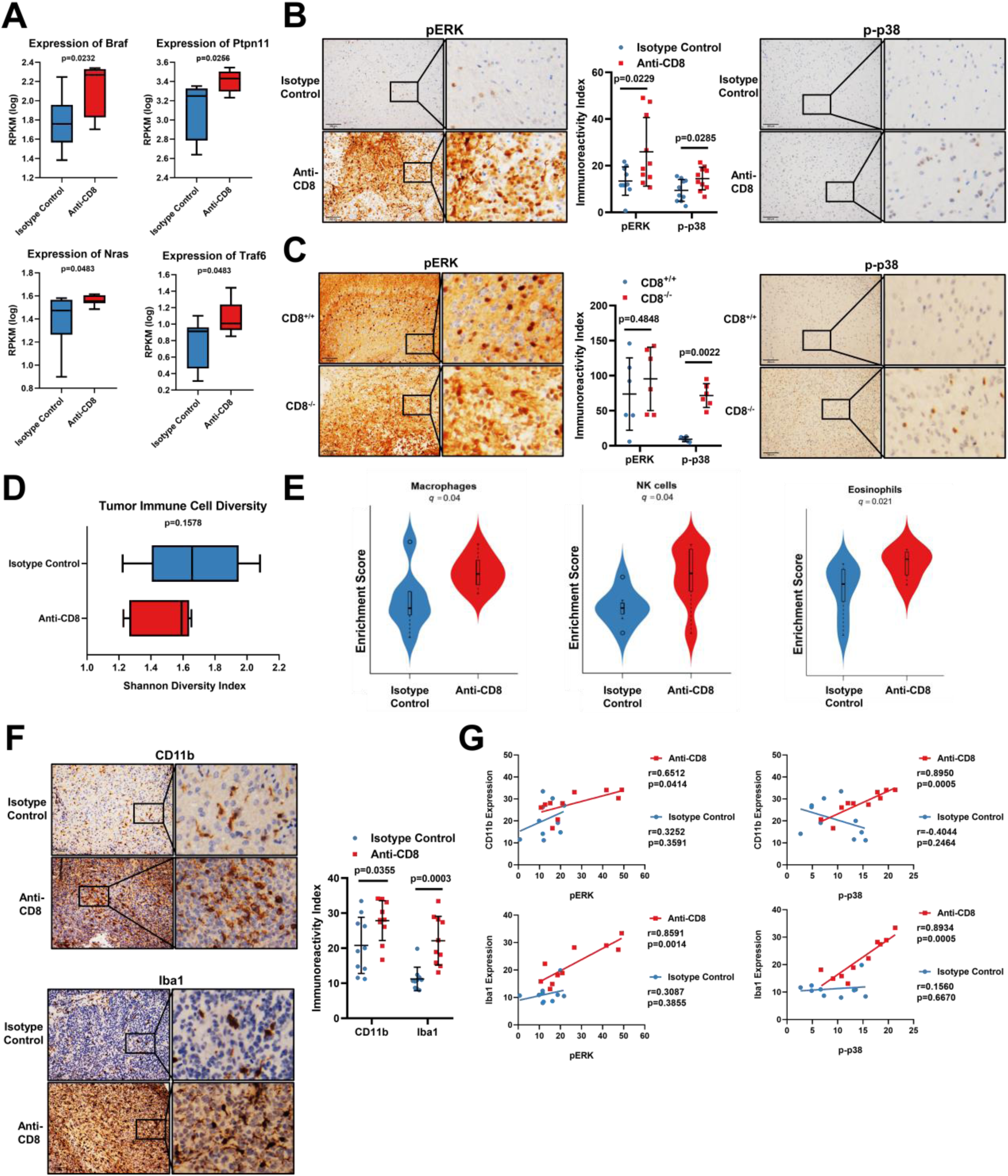
MAPK pathway (ERK and p38) activation is increased gliomas that develop in the absence of CD8+ T-cells and is correlated to increased macrophage/microglial infiltration. (A) *Braf, Ptpn11* (n=16; p=0.0256), *Nras*, and *Traf6* (n=16) expression compared between tumors from mice treated with anti-CD8 and isotype control antibodies by analysis of RNA-seq data in a Pten^-/-^ transgenic murine glioma model. (B) pERK immunoreactivity index compared between groups. Quantification of pERK and p-p38 (n=20). p-p38 immunoreactivity index compared between groups. (C) pERK immunoreactivity index compared between tumors from CD8^-/-^ and CD8^+/+^ mice by IHC in a Stat3^-/-^ transgenic murine glioma model. Quantification of pERK and p-p38 (n=16) expression. p-p38 compared groups. (D) Shannon diversity index of immune cell populations in tumors developed in the absence of CD8+ T-cells compared to the isotype control (n=20). (E) Macrophage, NK cell, and eosinophil (n=20) gene expression signature between groups by ciber-sorting of RNA-seq data. (G) CD11b immunoreactivity index compared between groups by IHC. Quantification of CD11b and Iba1 (n=20) expression. Iba1 expression compared between groups. (G) Correlation between CD11b expression and pERK (n=10). Correlation between CD11b expression and p-p38 (n=10). Correlation between Iba1 expression and pERK (n=10). Correlation between Iba1 expression and p-p38 (n=10).

The oncogenic MAPK signaling pathway has distinct branches that include the ERK, p38, and JNK effectors. We next investigated the expression of other known activators of these signaling cascades. *Nras* is an activator of the ERK cascade and was over-expressed in these gliomas (p=0.0483). *Traf6*, an activator of the p38 cascade, was also over-expressed (p=0.0483). There were no differences in expression of known activators of the JNK cascade (data not shown). Given the over-expression of these activators of MAPK signaling in gliomas that developed in the absence of CD8+ T-cells, we then investigated the functional activity of the ERK and p38 cascades. Gliomas that developed in the absence of CD8+ T-cells had elevated activation of these cascades as demonstrated by immunoreactivity for ERK and p38 phosphorylation (p). pERK in gliomas that developed in the absence of CD8+ T-cells revealed intense diffuse positivity with a mean immunoreactivity score of 25.91 as compared to gliomas that developed in the presence of CD8+ T-cells that had a mean immunoreactivity score of 13.41 (p=0.0229; **Fig. 3B**). p-p38 expression in gliomas that developed in the absence of CD8+ T-cells demonstrated scattered focal positivity with a mean immunoreactivity score of 14.50 as compared to gliomas that developed in the presence of CD8+ T-cells that had a mean immunoreactivity score of 9.41 (p=0.0285).

To determine if this oncogenic signaling was unique to the PDGF^+^Pten^-/-^ murine glioma model, we evaluated the phosphorylation of the ERK and p38 cascades in a PDGFB^+^RCAS^-^Stat3^-/-^ transgenic glioma model in which CD8+ T-cells were depleted by germline homozygous knock-out of CD8 [9]. In this model, pERK was elevated with a mean immunoreactivity score of 95.39 and 73.76 in the gliomas from either the CD8^-/-^ or the CD8^+/+^ background, respectively (p=0.4848; **Fig. 3C**). Similar to the PDGF^+^Pten^-/-^ murine glioma model, the PDGFB^+^RCAS^-^Stat3^+/+^ gliomas demonstrated similar findings in p-p38. More specifically, gliomas in the CD8^-/-^ background revealed scattered to moderate p-p38 positivity with a mean immunoreactivity score of 71.56 as compared to a mean immunoreactivity score of 9.18 in the CD8^+/+^ gliomas (p=0.0022).

We next investigated whether the expression of upstream regulators of the MAPK pathway correlated with phosphorylation of effectors in the PDGF^+^Pten^-/-^ murine glioma model. *Braf* (r=0.7276; p=0.0408) and *Ptpn11* (r=0.8499; p=0.0075) expression by RNA-seq correlated with pERK in gliomas that developed in the absence of CD8+ T-cells (**Supplementary Fig. S2A/B**). *Nras* expression did not correlate with pERK in either gliomas (**Supplementary Fig. S2C**). *Traf6* showed a trend in expression with p-p38 in gliomas that developed in the absence of CD8+ T-cells (r=0.6496; p=0.0813) (**Supplementary Fig. S2D**). Together, these results indicate that MAPK alterations are stochastic events that are suppressed by CD8+ T-cells during the progression of gliomas. Our results also confirm the integrity of the MAPK signaling cascade in these gliomas, as expression of upstream members of the pathway are associated with phosphorylation of the downstream effectors, which is indicative of activation.

### Macrophage/microglial infiltration is increased in gliomas that develop in the absence of CD8+ T-cells

We hypothesized that the immunogenicity of gliomas that developed in the absence of CD8+ T-cells relates to a distinct immune microenvironment. We determined the Shannon diversity index between the immune cell populations within glioma groups. Gliomas that developed in the absence of CD8+ T-cells had a collective index ranging from 1.228 to 1.652 while those that developed with intact immunity had an index ranging from 1.223 to 2.081 (p=0.1578; **Fig. 3D**) indicating a greater diversity of glioma infiltrating immune cell populations in mice that had CD8+ T-cells. We then performed ciber-sorting of the main immune cell populations by RNA-seq as previously done [15]. This analysis showed an increase in the expression signature of natural killer (NK) cells (q=0.04), macrophages (q=0.04), and eosinophils (q=0.021; **Fig. 3E**).

IHC staining confirmed that macrophage/microglia infiltration in gliomas was enhanced in the absence of CD8+ T-cells. CD11b is a marker of macrophages that has been reported to be associated with pro-inflammatory polarization [16]. CD11b expression revealed diffuse positivity with a mean immunoreactivity score of 27.87 in gliomas that developed in the absence of CD8+ T-cells as compared to gliomas with a mean immunoreactivity score of 20.79 that developed in the presence of CD8+ T-cells (p=0.0355; **Fig. 3F**). Another marker for macrophages, Iba1, also showed higher expression with a mean immunoreactivity score of 22.12 in gliomas that developed in the absence of CD8+ T-cells relative to a mean immunoreactivity score of 11.22 in gliomas that developed in the presence of CD8+ T cells (p=0.0003). These results indicate that macrophage/microglial infiltration is increased in gliomas that developed in the absence of CD8+ T-cells.

### MAPK signaling is correlated with macrophage infiltration in gliomas that develop in the absence of CD8+ T-cells

Since we encountered robust yet variable macrophage infiltration and MAPK pathway activation in the gliomas that developed in the absence of CD8+ T-cells, we investigated whether this signaling correlated with the degree of immune infiltration. CD11b expression correlated with pERK in gliomas that developed in the absence of CD8+ T-cells (r=0.6512; p=0.0414) but not in those gliomas that developed in the presence of CD8+ T-cells (r=0.3252; p=0.3591; **Fig. 3G**). CD11b also correlated with p-p38 in gliomas that developed in the absence of CD8+ T-cells (r=0.8950; p=0.0005) but not in gliomas developed with CD8+ T-cells present (r=0.4044; p=0.2464). Similarly, Iba1 expression correlated with pERK in gliomas that developed in the absence of CD8+ T-cells (r=0.8591; p=0.0014) but not in gliomas that developed in the presence of CD8+ T cells (r=0.3087; p=0.3855). Iba1 also correlated with p-p38 in gliomas that developed in the absence of CD8+ T-cells (r=0.8934; p=0.0005) but not in gliomas with intact immunity (r=0.1560; p=0.6670).

We then investigated the relationship between MAPK signaling and macrophage/microglial infiltration in human GBM. Messenger RNA (mRNA) expression analysis of 161 GBM patients in the Tumor Cancer Genome Atlas (TCGA) revealed a correlation between ITGAM (CD11b) and *MAPK13* (p38delta), an activator of the p38 cascade (r=0.6194; p<0.0001; **Supplementary Fig. S3A**). mRNA expression analysis of GBM patients in the TCGA revealed a correlation between AIF1 (Iba1) and *RRAS*, an activator of the ERK cascade (r=0.6141; p<0.0001; **Supplementary Fig. S3B**). AIF1 levels also correlated with *MAPK13* (r=0.6008; p<0.0001: **Supplementary Fig. S3C**), and *MAP3K8*, an activator of the ERK cascade (r=0.8159; p<0.0001; **Supplementary Fig. S3D**). Thus, MAPK activation and macrophage/microglial recruitment were correlated in both murine gliomas and human GBM patients. This association appears to be strengthened by the absence of CD8+ T-cells during glioma development.

### Gliomas that developed in the absence of CD8+ T-cells exhibit a gene signature associated with pro-inflammatory macrophage polarization

Given that macrophage/microglial infiltration was robust in gliomas that developed in the absence of CD8+ T-cells, we explored whether this immune cell infiltration was associated with an inflammatory milieu. Genes of proteins known to interact with ITGAM (CD11b) [17] showed a trend for elevated mRNA expression in gliomas that developed in the absence of CD8+ T-cells (ANOVA p=0.0693; **Fig. 4A**). The expression of genes known to interact AIF1 (Iba1) [17] were no different between the groups of mice (ANOVA p=0.5421; **Fig. 4B**). As macrophages can exist either in a non-polarized state or along a continuum that is pro- or anti-inflammatory [18], we evaluated such phenotypic changes in these gliomas. Gliomas that developed in the absence of CD8+ T-cells had an increased pro-inflammatory phenotype by a gene signature that is reported to contribute to such a polarization (ANOVA p=0.0003; **Fig. 4C**) [19, 20]. A gene signature demonstrated to contribute to an anti-inflammatory macrophage polarization [19] did not reveal significant differences (data not shown). Gene set enrichment analysis (GSEA) revealed an increased pro-inflammatory macrophage phenotype that has been described by curated datasets in the gliomas that developed in the absence of CD8+ T-cells, with a normalized enrichment score (NES) of 1.2797 compared to those that developed in the presence of CD8+ T cells (FDR=0.1431). Together, these results indicate that gliomas that developed in the absence of CD8+ T-cells exhibit a gene signature associated with pro-inflammatory macrophage polarization.

**Figure 4:**
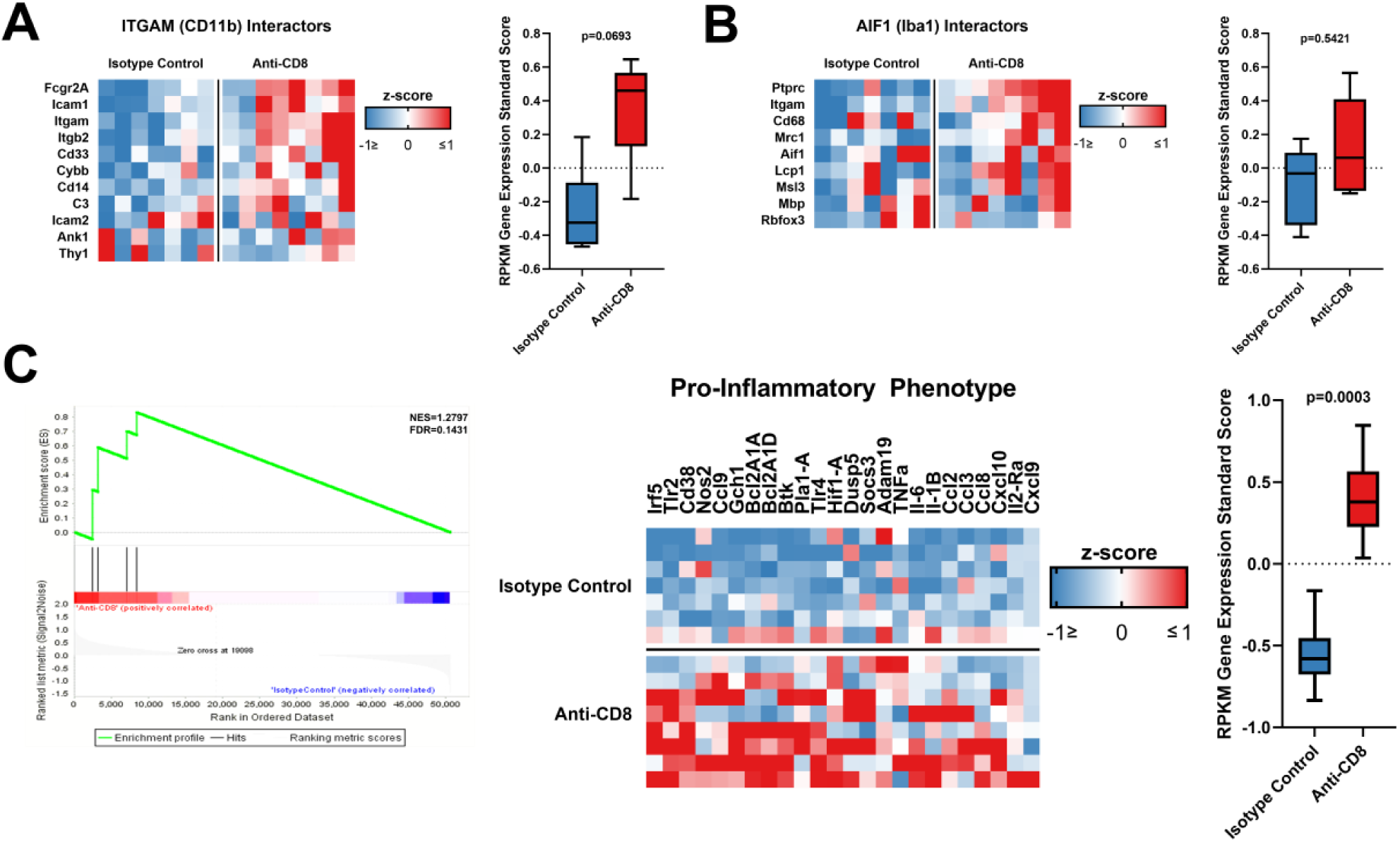
Macrophages/microglia in tumors that develop in the absence of CD8+ T-cells have an enriched pro-inflammatory phenotype. RNA-seq based expression analysis of genes of proteins known to interact with ITGAM (CD11b) (A) and of genes of proteins known to interact with AIF1 (Iba1) (B) in tumors developed in the absence of CD8+ T-cells versus the isotype control (n=15). (C) The expression signature of pro-inflammatory phenotype in macrophages/microglia is compared between tumors developed in the absence of CD8+ T-cells and the isotype control by RPKM z-score (n=15) using the GSEA approach (left, NES=1.2797; FDR=0.1431), heatmap (middle), and box-plot (right) (n=15). p value calculated using ANOVA test.

## Discussion

Cancer immunoediting shapes tumor evolution by eliminating immunogenic tumor cell variants that can evade immune surveillance while others are selected for resistance to immune recognition [21]. Whereas this concept has previously been suggested in GBM [22, 23], our study directly investigated the effects of CD8+ T-cell-dependent immunoediting in the development of gliomas. We obtained experimental evidence of immunoediting by CD8+ T-cells in a transgenic murine glioma model as depletion of this immune cell population had a profound effect on tumor evolution.

We sought to explore genomic differences as a result of tumorigenesis in the absence of CD8+ T-cells. Low mutational burden is a common explanation for the relative lack of immunogenicity in GBM [24]. We found no change in the number of point mutations as a result of the absence of T-cells. Our results imply that CD8+ T-cells do not protect against point mutations or influence this facet of immunogenicity to an appreciable extent. Yet, it is important to consider that in this particular glioma model, mice may not survive long enough for the development and selection of mutations other than those with a strong advantage for cell growth (i.e. Trp53).

In the absence of CD8+ T-cells, we observed an increase in the frequency of glioma gene fusions. Gene fusions are known to be strong drivers of neoplasia as well as the formation of neoantigens from junctional epitopes [25]. Bioinformatic modeling predicted these gene fusions to have high binding affinity to MHC-I. Whereas the antigenic properties predicted warrant experimental validation, this finding is interesting as junctional epitopes from gene fusions are a potential source for development of neoantigen-based vaccination.

Genomic instability was also observed in gliomas developed in the absence of CD8+ T-cells. In our study, such instability manifested as aneuploidy. Previous research correlated aneuploidy with immune evasion in cancer, including GBM [10]. Meanwhile, we show that gliomas developed in the absence of CD8+ T-cells have both, high aneuploidy and a more immunogenic phenotype, reinforcing the biological importance of genomic instability in cancer [26]. Chromosomal instability drives innate immune activation via cGAS-STING in cancer [12], and in support of this, we showed upregulation of genes related to this pathway in gliomas that developed in the absence of CD8+ T-cells [13].

Surprisingly, whereas depletion of CD8+ T-cells during glioma development had profound effects on tumor genotype, phenotype, immunogenicity and microenvironment, depletion of these immune effector cells did not lead to shorter survival. CD8+ T-cell depletion did not alter survival of other transgenic glioma models in which tumors develop de novo [9]. Previous studies also showed that whereas immunosuppression can increase the incidence of cancer, it usually does not affect survival upon cancer diagnosis [27]. These observations in the face of tumor progression is termed the “Hellstrom paradox” [28, 29]. How this paradoxical situation manifests remains unresolved.

We recently reported that the presence of MAPK activating mutations in *BRAF* and *PTPN11* in recurrent GBM patients who responded to PD-1 blockade [30]. Given the implication of these genes and MAPK in response to immunotherapy, we explored how this pathway is affected by the presence of CD8+ T-cells during glioma development. Whereas these genes were not mutated in the absence of CD8+ T-cells, we did find them over-expressed in this setting, along with several other known activators of ERK and p38 signaling [30]. In PDGFB^+^RCAS^-^Stat3^-/-^ gliomas, a separate transgenic murine glioma model, the p38 cascade was more activated in the absence of CD8+ T-cells but not in the ERK cascade. MAPK activation has been extensively studied as a driver of innate immunity with effector pro-inflammatory polarization [31]. Our results indicate that CD8+ T-cells protect against such oncogenic signaling that albeit was associated with increased response to PD-1 blockade therapy in this disease [30].

We observed increased infiltration of macrophages/microglia in gliomas that developed in the absence of CD8+ T-cells. Our analysis revealed a pro-inflammatory phenotype that was observed in gliomas that developed in the absence of these immune cells. Future studies will require investigation as to their role in this context. We did encounter a strong relationship between oncogenic MAPK signaling and macrophage/microglial recruitment in these gliomas. mRNA expression analysis of human GBM patients *in silico* revealed a strong correlation between levels of known activators of the MAPK signaling pathway and macrophage markers, asserting the biological significance of these findings in the human disease. Recently, it was demonstrated that osteopontin which has been associated with an immunosuppressive phenotype, is expressed by gliomas and is a strong chemotactic for macrophage infiltration that is associated with the p38 MAPK signaling pathway [32, 33]. On the other hand, MAPK is known to promote a pro-inflammatory innate immune response [31]. Consistent with this, we established an association between MAPK pathway activation and pro-inflammatory macrophage recruitment in gliomas. Whereas we observed that CD8+ T-cells dampen MAPK signaling in gliomas, the effect of MAPK signaling on pro-inflammatory microenvironment in the context of glioma immunoediting remains to be elucidated. Another question that remains unanswered is whether MAPK signaling in tumor cells versus macrophages/microglia has a different effect on the microenvironment.

CD8+ T-cell recognition of tumor antigens drives the immunological sculpting of a developing cancer [4]. Our study suggests that glioma immunoediting is the consequence of a CD8+ T-cell-dependent selection process leading to immune evasion. While we found strong differences between gliomas that developed with or without this immune cell population, we also found important variation of these features within the group of gliomas that developed in the absence of CD8+ T-cells. This indicates that glioma evolution is stochastic, with recurrent events that can lead to immune recognition or evasion. Cancer immunoediting mediated by CD8+ T-cells simultaneously protects against certain evolutionary hallmarks of cancer and selects for tumor cell variants that are capable of immune evasion. Understanding the interplay between glioma progression and anti-tumoral immunity is essential for the development of effective immunotherapy for this disease. Manipulation of tumor evolution towards a more immunogenic path may be explored by surpassing the CD8+ T-cell-mediated immune evasion in this disease.

## Methods

### In vivo models

PDGF^+^*PTEN*^-/-^ gliomas were generated by intracranial injection of a PDGF-IRES-Cre retrovirus into the subcortical white matter of adult mice with the *Pten*^lox/lox^ genotype (C57BL/6 background, Jackson Lab Stock No: 006440). Gliomas generated in hosts with intact immunity can graft and grow in syngeneic immune-competent recipients. Transplantation was done by extracting the gliomas that developed in the absence of CD8+ T-cells that were dissociated with 2.5% TrypLE in PBS upon which it was cultured with DMEM + 0.5% fetal bovine serum, 1% N2, 1% penicillin/streptomycin, 10ng/mL PDGF-AA, and 10ng/mL FGF (Basal FGF PDGF), and then intracranially implanted into both groups of animals. Animals were anesthetized with ketamine/xylazine (115/17 mg/kg) and a burr hole was made through stereotactic injection using a 10 μL Hamilton syringe (Hamilton; Reno, NV) with a 30-gauge needle while animals were mounted on a stereotactic apparatus (Harvard Apparatus; Holliston, MA). For survival analysis, animals losing ≥ 30% of their body weight or having trouble ambulating, feeding, or grooming were euthanized by CO_2_ administration followed by cervical dislocation. After conclusion of survival analysis, animals were sacrificed. Brains were harvested, sectioned, and stained by immunohistochemistry (IHC), and hematoxylin & eosin (H&E).

### CD8+ T-cell depletion

Animals were treated with anti-mouse CD8α (BE0004-1, BioXCell) and an anti-rat IgG2a isotype control, anti-trinitrophenol (BE0089, BioXCell), one week prior to glioma induction at 0.2 mg i.p. and continued twice per week in order to achieve consistent CD8+ T-cell depletion. Depletion was monitored at three time points starting from intracranial virus injection, 3 weeks apart.

### Immunohistochemistry

Formalin-fixed, paraffin-embedded (FFPE) material from animals was deparaffinized with xylene and antigen retrieval was performed with 10mM sodium citrate buffer (pH=6). IHC was performed with a 1:150 diluted rat monoclonal antibody against MHC-I (AB15680) (Abcam; Cambridge, UK), 1:150 diluted rabbit polyclonal antibody against MHC-II (AB180779) (Abcam; Cambridge, UK), 1:2000 diluted rabbit monoclonal antibody against CD11b (AB133357) (Abcam; Cambridge, UK), 1:3000 diluted rabbit monoclonal antibody against Iba1 (AB178846) (Abcam; Cambridge, UK), 1:250 diluted rabbit monoclonal antibody against p38 (4511) (Cell Signaling; Danvers, MA), and 1:250 diluted rabbit monoclonal antibody against pERK (4370) (Cell Signaling; Danvers, MA). Counterstaining was performed on the same material with hematoxylin (Abcam; Cambridge, UK). IHC staining was performed on a Leica Bond-Max automatic immunostainer (Bannockburn, IL).

Machine learning of IHC quantification was used by setting parameters to identify positivity and subtracting counter-stained/negative background using ImageJ software (available from the National Institute of Health). The substrate optical density and intensity was determined by quantifying the positive nuclei or cytoplasm (dependent on particular antibody assessed) paired with the intensity of the staining in neoplastic tumor cell regions of the brain at 4x light microscopic magnification in order to infer immunoreactivity index.

### Bioanalysis of genomic instability

Mutational burden was called by analysis of germline and somatic variation by Strelka2 Small Variant Caller (Illumina; San Diego, CA). The germline identification employed a tiered haplotype model adaptively selected between assembly and alignment-based haplotyping at each variant locus. The aneuploidy score from somatic copy number alterations was calculated by calling the presence or absence of amplifications or deletions as described in practice [10]. Aneuploidy score as a comparison between two groups was called by copy number detection, implemented in the software package CNVkit, using both targeted reads and non-specifically captured off-target reads to infer copy number evenly across the genome [11]. STAR-Fusion was used to identify candidate fusion events [34]. The impact of the fusion event on coding regions was explored by invoking the ‘examine_coding_effect’ parameter. Only the candidates that affected the coding regions were retained for fusion load analysis. Copy number variance events were counted and the percentage of the genome affected by such events was calculated as described [10].

### Protein interaction analysis

Evidence-based protein-protein interaction networks were evaluated via the STRING database (http://string-db.org). STRING provided hierarchical and self-consistent orthology annotations for all interacting proteins at various levels of phylogenetic resolution in order to assess interaction probability with a given gene. Analysis included genes with function gene ontology with a predicted interaction probability greater than 0.73.

### Calculation of Shannon index

For each tumor, the Shannon diversity was estimated using the command ‘entropy.empirical’ from the ‘entropy’ R package. This was calculated on the basis of the estimated prevalence of different immune cell compartments found in the tumor. The Shannon diversity score, H, followed the formula H = −Σpi × log(pi).

### Prediction of neoantigen binders

Novel 9–11mer peptides that could arise from identified non-silent mutations or gene fusions present in the tumor samples were determined. We used the pVAC-Seq54 pipeline with the NetMHCcons55 binding strength predictor to identify neoantigens [35, 36]. The predicted IC50 inhibitory concentration binding affinities and rank percentage scores were calculated for all peptides that bound to each of the tumor’s HLA alleles. Using established thresholds, predicted binders were considered to be those peptides that had a predicted binding affinity < 500 nM.

### Statistical analysis

All statistical analysis was conducted using GraphPad Prism 8 (GraphPad Software; San Diego CA) and R version 3.1.2 (R Core Team; Vienna, Austria). Numerical data was reported as mean ± SEM. Mann-Whitney or unpaired Student’s t test was used for two group comparisons. ANOVA was used for more than two groups. Kaplan-Meier survival curves were generated and a log rank test was utilized to compare survival distribution. Mutational and gene fusion counts were analyzed using a binomial model for count data with over-dispersion. All reported q values were calculated by statistical adjustment of p values using the Benjamini-Hochberg constant. The Pearson coefficient of correlation (r) was calculated by the linear regression of the data points according to the goodness of fit where significance was determined at which point the slope was significantly non-zero. *In silico* analysis was performed with 161 cataloged, human GBM samples provided by the Tumor Cancer Genome Atlas (TCGA) with mRNA expression data available (RNAseq V2 RSEM).

## Acknowledgments

This work was funded by 5DP5OD021356-05 (AMS), P50CA221747 SPORE for Translational Approaches to Brain Cancer (AMS), developmental funds from the Robert H. Lurie Cancer Center Support Grant #P30CA060553 (AMS), and the Matthew Larson Foundation IronMatt Research Award (AMS). Histology services were provided by the Northwestern University Mouse Histology and Phenotyping Laboratory which is supported by NCI P30-CA060553 awarded to the Robert H. Lurie Comprehensive Cancer Center.

---

**Supplementary Figure S1**: **Transgenic murine glioma model possesses increased CD4+ and CD8+ T-cell infiltration early in tumor development**. (A) Representative bioluminescence activity of tumor captured 21 and 35 days post-tumor induction. *Ex vivo* flow cytometry of the brain tumors are shown in the right panel. (B) Bioluminescence was measured at seven time points between 21 and 63 days post-tumor induction (n=8/time point). (C) Kaplan-Meier survival curve illustrates the time points of flow cytometric analysis relative to initiation of tumor-related death. (D) CD4+ and CD8+ T-cell tumor infiltration normalized against the contralateral brain hemisphere, and bioluminescence signal at 21 and at 35 days post-tumor induction with provided standard deviation (n=16).

**Supplementary Figure S2**: **Expression of upstream activators of the MAPK pathway correlates with functional activation of the ERK and p38 cascades**. (A) Correlation between *Braf* expression and pERK in tumors from mice treated with anti-CD8 (n=8) and isotype control (n=8) antibodies. (B) Correlation between *Ptpn11* expression and pERK in tumors from mice treated with anti-CD8 (n=8) and isotype control (n=8) antibodies. (C) Correlation between *Nras* expression and pERK in tumors from mice treated with anti-CD8 (n=8) and isotype control (n=8) antibodies. (D) Correlation between *Traf6* expression and p-p38 in tumors from mice treated with anti-CD8 (n=8) and isotype control (n=8) antibodies.

**Supplementary Figure S3**: **Known activators of MAPK signaling correlate to macrophage markers in human GBM patients**. (A) Correlation of mRNA expression analysis of 161 GBM patients in the TCGA library between ITGAM (CD11b) and *MAPK13* (n=161). (B) Correlation between AIF1 (Iba1) and *RRAS* (n=161). (C) Correlation between AIF1 and *MAPK13* (n=161). (D) Correlation between AIF1 and *MAP3K8* (n=161).

