## Supplementary figures and images for "CD8+ T-cell-mediated immunoediting influences genomic evolution and immune evasion in murine gliomas"

### Supplementary Figures

#### Supplementary Figure 1

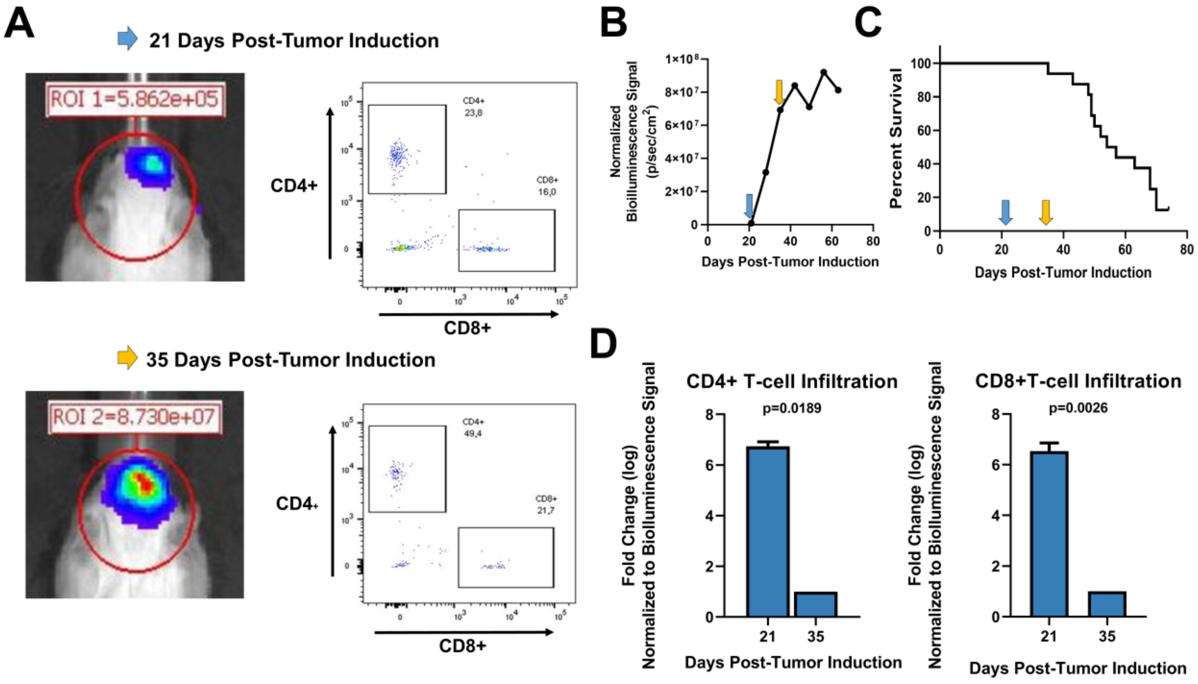

#### Supplementary Figure 2

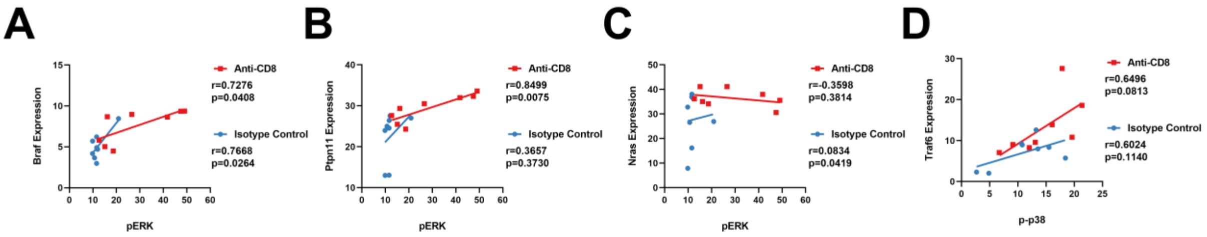

#### Supplementary Figure 3

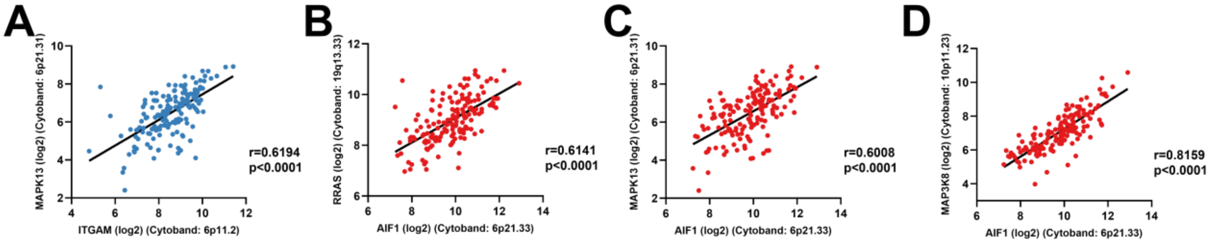
